# Automated atlas-based multi-label fetal cardiac vessel segmentation in Congenital Heart Disease

**DOI:** 10.1101/2022.01.14.476320

**Authors:** Paula Ramirez Gilliland, Alena Uus, Milou P.M. van Poppel, Irina Grigorescu, Johannes K. Steinweg, David F.A. Lloyd, Kuberan Pushparajah, Andrew P. King, Maria Deprez

**Affiliations:** School of Biomedical Engineering and Imaging Sciences, King’s College London, London, UK

**Keywords:** Automated Segmentation, Congenital Heart Disease, Fetal MRI, Fetal Cardiac Imaging, Label Propagation

## Abstract

Congenital heart disease (CHD) is the most commonly diagnosed birth defect. T2w black blood MRI provides optimal vessel visualisation, aiding prenatal CHD diagnosis. Common clinical practice involves manual segmentation of fetal heart and vessels for visualisation and reporting purposes.

We propose an automated multi-label fetal cardiac vessels deep learning segmentation approach for T2w black blood MRI. Our network is trained using single-label manual segmentations obtained through current clinical practice, combined with a multi-label anatomical atlas with desired multi-label segmentation protocol. Our framework combines deep learning label propagation with 3D residual U-Net segmentation to produce high-quality multi-label output well adapted to the individual subject anatomy.

We train and evaluate the network using forty fetal subjects with suspected coarctation of the aorta, achieving a dice score of 0.79 ± 0.02 for the fetal cardiac vessels region. The proposed network outperforms the label propagation and achieves a statistically equivalent performance to a 3D residual U-Net trained exclusively on manual single-label data (p-value*>*0.05). This multi-label framework therefore represents an advancement over the single-label approach, providing label-specific anatomical information, particularly useful for assessing specific anomaly areas in CHD.

## 1. Introduction

Congenital heart disease (CHD) encompasses a set of cardiac malformations present from birth, with varying degrees of severity and anatomical variability (Liu et al., 2019). CHD is the most frequently diagnosed congenital condition, and the dominant cause of congenital malformation-related deaths (Mendis et al., 2011).

Fetal cardiac MR (CMR) has proven its efficacy as an adjunct to echocardiography for detecting CHD prenatally (Dong and Zhu, 2018; Salehi et al., 2021), with T2w black blood MRI being particularly advantageous for vascular assessments (Lloyd et al., 2016, 2019).

Routine clinical practice assessments involve manual segmentation of ROIs, which is laborious and time consuming. Hence, there is an important clinical need for automated fetal CMR segmentation methods, with deep learning being the method of choice for state-of-the-art automated segmentation approaches (Hesamian et al., 2019) and adult CMR segmentation (Chen et al., 2020).

Training a CNN for segmentation normally requires a large number of manually labelled examples. In this paper we propose a novel deep learning framework that combines label propagation from a multi-label atlas with manual single-label vascular ROI segmentations (see Fig. 1) for training of a 3D residual U-Net to predict high quality multi-label segmentations, well adapted to the individual subject anatomy.

**Figure 1:**
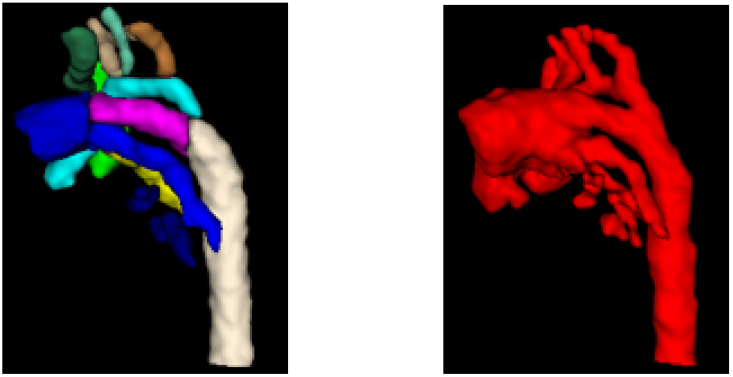
3D models of the two types of segmentation used: a multi-label cardiac atlas (left), and single-label manual segmentations (right).

We show that this approach is able to propagate the multi-label protocol from the fetal cardiac atlas while achieving high accuracy in individual images driven by the manual single cardiac vessel labels. These highly accurate fully automated multi-label segmentations will aid visualisation of the fetal cardiac vessels for prenatal diagnostic reporting purposes, and provide the basis for automated vessel biometry and detection.

## 2. Related Work

### 2.1. Deep learning segmentation

U-Net (Ronneberger et al., 2015) is one of the most widely used deep neural network architectures, due to its performance, generalisability and efficiency (Isensee et al., 2018). U-Net based architectures have been successfully employed for adult cardiac CMR localisation (Payer et al., 2017), and segmentation in CHD (Xu et al., 2019).

Residual U-Net (Zhang et al., 2018) combines the advantages of Residual connections (He et al., 2016) with U-Net, to produce an enhanced network that eases training and information propagation without degradation. This type of architecture has demonstrated high segmentation performance on adult cardiac segmentation tasks (Kerfoot et al., 2018; Vesal et al., 2020). While residual U-Net can produce highly accurate automated segmentation in individual images, it is restricted to the protocol of the available manual labels, which in our case is a single-label vessels ROI segmentation (Fig. 2 left).

**Figure 2:**
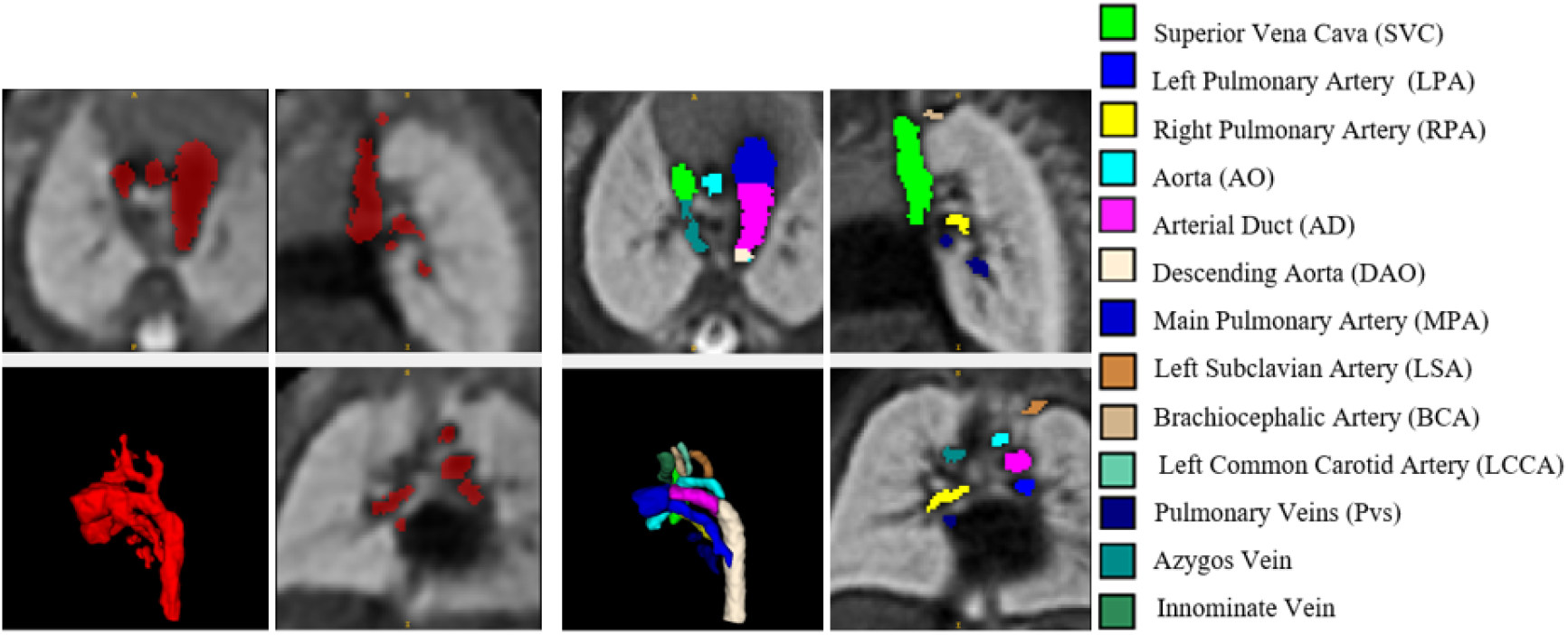
Left: 3D SVR reconstructed image of one of the subjects with corresponding single manual label. Right: 3D T2w MRI CoA atlas with corresponding multilabel vessel segmentations.

### 2.2. Label propagation

In atlas-based label propagation, the label information from a given atlas is transferred to an individual subject via image registration (Heckemann et al., 2006). With deep learning becoming an increasingly popular choice for image registration due to high computational efficiency and optimal performance (Tustison et al., 2019), we propose the use of VoxelMorph (Balakrishnan et al., 2019) for label propagation from a multi-label atlas (Fig. 2 right) to individual subjects. While label propagation can transfer any desired segmentation protocol from the atlas to the individual images, its accuracy is limited by the registration quality, and thus residual U-Net may produce more accurate results.

### 2.3. Automated fetal segmentation

In fetal MRI segmentation, U-Net based architectures have been employed for fetal brain, eye and thorax segmentation (Payette et al., 2020; Salehi et al., 2018; Uus et al., 2021b,a). The only automated fetal cardiac segmentation method in literature consists of a random forest for global cardiac, liver and lungs detection (Keraudren et al., 2015). To our knowledge, the present paper proposes the first multi-label segmentation of fetal cardiac structures, aimed at patients with CHD.

### 2.4. Contribution

Our deep learning framework combines advantages of highly accurate residual U-Net segmentation with deep learning label propagation of any desired protocol, fine-tuned by the available single-label manual segmentations (see Fig. 3). We show that the resulting multilabel residual U-Net segmentation outperforms the label propagation, while achieving the same accuracy as the single-label residual U-Net trained on manual labels.

**Figure 3:**
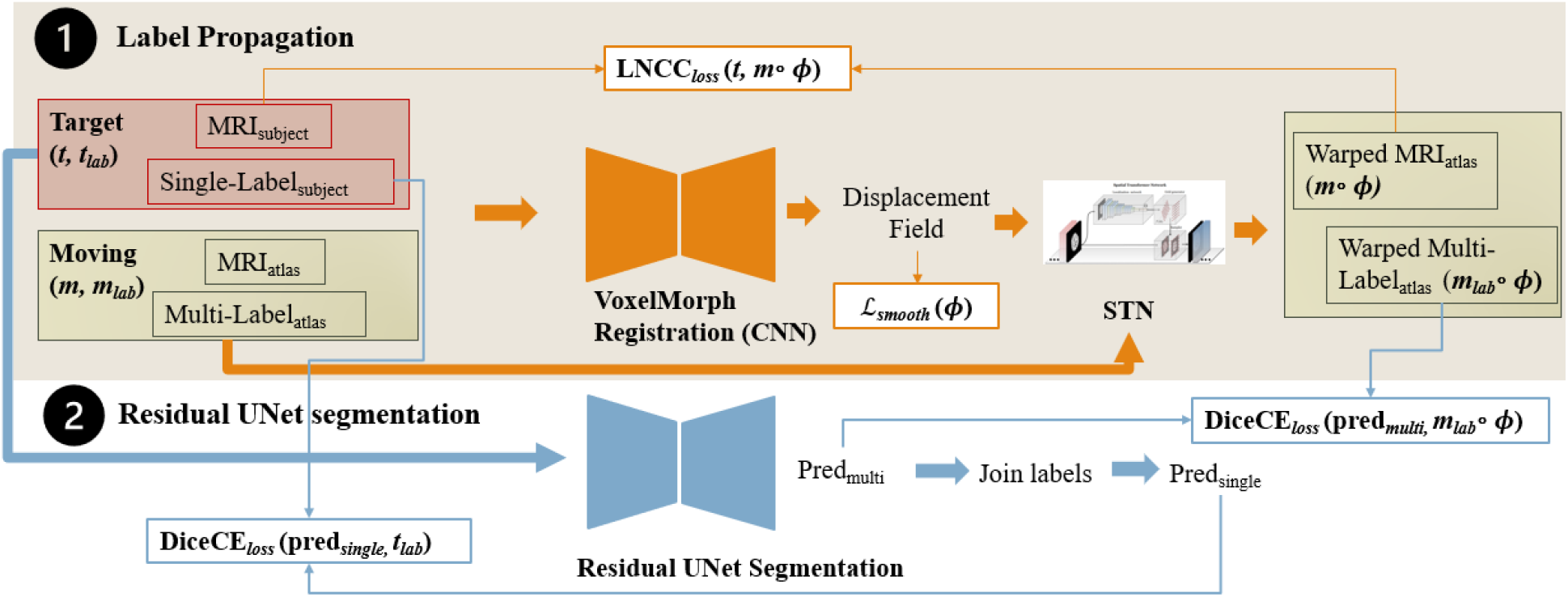
Overall architecture presented. The top half and orange arrows depict the label propagation framework, while the lower half and blue arrows illustrate the segmentation pipeline. Arrows of increased thickness depict the various stages *t* and *m* go through, while thinner arrows indicate the volumes used in the losses (detailed in subsequent sections).

## 3. Methods

### 3.1. Data specifications

Our dataset consists of 40 fetal subjects with suspected coarctation of the aorta (CoA), 32 weeks mean GA. The datasets were acquired at Evelina London Children’s Hospital using a 1.5 T Tesla Ingenia MRI system, T2-weighted SSFSE sequence (repetition time = 20,000 ms, echo time = 50ms, flip angle = 90°, voxel size = 1.25 × 1.25 mm, slice thickness = 2.5 mm and slice overlap = 1.25 mm). The raw datasets comprised 6-12 multi-slice 2D stacks, covering the fetal thorax in three orthogonal planes, and were reconstructed using Slice-to-Volume Registration (SVR) (Kuklisova-Murgasova et al., 2012; Kainz et al., 2015) to 0.75 mm isotropic resolution. Each subject image was manually segmented by trained clinicians using ITK-SNAP (Yushkevich et al., 2006), encompassing the main cardiac vessels region (single label).

We employ a 3D CoA cardiac atlas^1^ from which we select 13 manually segmented vascular regions, including two vessels (innominate vein and azygos vein) which are not present in the manual segmentations. We crop all data to the cardiac vessels region (see Fig. 2).

We split the subjects into a training set (n=32), validation set (n=3) and testing set (n=5), and normalise and rescale the intensity between 0 and 1 for network training.

### 3.2. Overall Architecture

The overall framework consists of two main steps: label propagation and automated segmentation (see Fig. 3). First, the label propagation network (VoxelMorph) is fully trained. The resulting propagated multi-class labels are then used to train a residual U-Net on the task of automated multi-label segmentation.

### 3.3. Label Propagation

#### 3.3.1. Registration network

We employ VoxelMorph (Balakrishnan et al., 2019) for registration-based label propagation. VoxelMorph is a CNN-based registration method, where the function mapping the input image pairs (atlas and subject image) to the aligning deformation field is learnt. VoxelMorph optimises the deformation field *ϕ* by considering a global function composed of shared parameters of a dataset of volume pairs, benefiting from high computational efficiency. This global function is parametrised via the kernels of the convolutional layers of a CNN. A Spatial Transformer Network (STN) (Jaderberg et al., 2015) then uses the resultant deformation field to warp the moving image. We use the subject images as target images (*t*), and the atlas as moving image (*m*).

#### 3.3.2. Label propagation loss functions

**We use Local Normalised Cross Correlation loss (LNCC**_*loss*_) (Balakrishnan et al., 2019) as a similarity loss function:

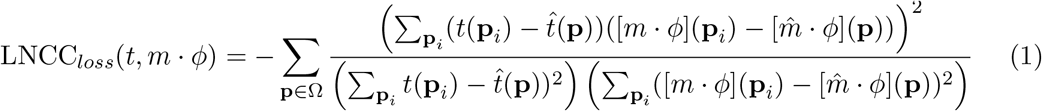

Here Ω is the target and moving image spatial domain; 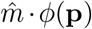 and 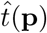 are the local mean intensities of an *n*^3^ volume of moving and target image, respectively; and **p***i* are the voxels contained within this *n*^3^ volume, which are iterated through.

We also include a **smoothness loss** (*ℒ*_*smooth*_) to encourage smoothness of the displacement field, thus ensuring the registration is physically realistic:

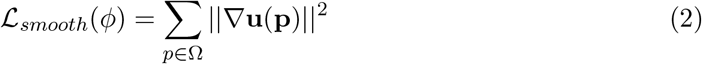

The spatial gradients are approximated via differences between neighbouring voxels (Balakrishnan et al., 2019).

We explored the use of an additional auxiliary segmentation loss between joined atlas labels and manual single-labels, however this resulted in insignificant registration overlap improvement (p-value>0.05).

Thus, in summary, **the total registration loss** *ℒ*_*reg*_ may be expressed as

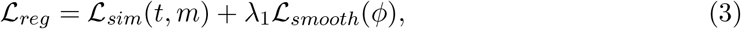

where *λ*_1_ is the smoothing loss weight.

#### 3.3.3. Registration network implementation details

We employ a U-Net based encoder-decoder architecture with skip connections for registration. Output channels are 16, 32, 32, 64 for the encoder (blocks of 3D strided convolutions with leaky ReLU activations); and 32, 32, 32 for the decoder (blocks of strided transpose 3D convolutions and leaky ReLU activations). This is followed by two convolutional blocks. See Appendix Section A for further details.

We train VoxelMorph registration network for 64,000 iterations, using a linearly decaying learning rate initialised at 5 × 10^*-*4^, and an Adam optimiser (default parameters *β*_1_ = 0.9, *β*_2_ = 0.999), with a weight decay of 1 × 10^*-*5^. We implement the LNCC_*loss*_ using *n* = 9, and *λ*_1_ = 0.2 for *L*_*smooth*_(*ϕ*), following hyperparameter tuning. We set the standard deviation of the smoothing kernel (applied to the velocity field) to 2.

We affinely register all subject images to the atlas prior to training. We use TorchIO (Pérez-García et al., 2021) data augmentation during training (random motion, spike, bias, blurring, ghosting, gamma and noise).

### 3.4. Multi-label segmentation

#### 3.4.1. Segmentation network

We use a 3D Residual U-Net (Zhang et al., 2018) for automated segmentation, with five encoder-decoder blocks (output channels 32, 64, 128, 256 and 512). We use Project MONAI implementation^2^, with convolution and upsampling kernel size of 3, two residual unit convolutions, PReLU activation and instance normalisation, and a dropout ratio of 0.5.

We employ an AdamW optimiser (Loshchilov and Hutter, 2017) with learning rate of 1 × 10^*-*4^, default *β* parameters and weight decay of 1 × 10^*-*5^. We train the network for 50,000 iterations, using intensity and spatial data augmentation techniques from Project MONAI (noise, smoothing, bias field, affine deformations, intensity shifts).

#### 3.4.2. Segmentation loss function

We train the segmentation network by combination of two losses. The first loss aims to minimise the difference between individual manual single-label segmentations and predicted multi-label segmentations, where the multi-class labels are joined into a single label. The second loss minimises the difference between predicted and propagated multi-label segmentations. The overall **segmentation loss** function *ℒ*_*seg*_ is expressed as

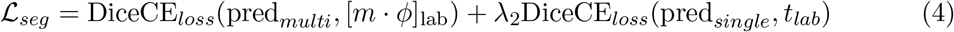

where pred*multi* are the multi-label residual U-Net predictions, pred*single* are single-label predictions (multi-label output labels joined together), *λ*_2_ is the single-class loss weight, [*m · ϕ*]_lab_ are the propagated labels from the atlas, and *t*_*lab*_ are the manual labels (single). We use the **soft dice and cross entropy loss** (Hatamizadeh et al., 2021) expressed as

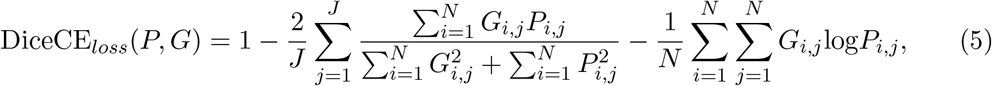

where *P* is the probabilistic prediction, *G* is the ground truth, *N* are the number of voxels, and *J* the number of classes. Refer to Fig. 3 for the summarised architecture with all losses.

### 3.5. Training experiments

Our experiments aim to quantitatively compare our proposed combined segmentation technique (*U-Net LP+manual*) with label propagation (*LP*), U-Net trained on propagated labels (*U-Net LP*) and U-Net trained on single-label manual segmentations (*U-Net manual*). Our training experiments are summarised as

- *LP* : Label Propagation using VoxelMorph.
- *U-Net LP* : Residual U-Net trained with propagated labels (*λ*_2_ = 0).
- *U-Net manual* : Residual U-Net trained with manual single-labels (*λ*_2_ = *∞*).
- *U-Net LP+manual* : Residual U-Net trained with both manual single-labels and propagated labels. We first train *U-Net LP+manual* using *λ*_2_ = 0.9 for 50,000 iterations, to ensure accurate multi-label classification, followed by 1,200 iterations with *λ*_2_ = 5.

This is to refine the segmentation of the vessels ROI, reducing any *LP* -induced biases.

## 4. Results

### 4.1. Quantitative evaluation

Table 1 includes results for mean Dice scores, 95% of Hausdorff Distance (HD95), and average surface distance on test set (n=5); comparing network predictions to manual single-labels (multi-class predictions are joined to form a single label).

**Table 1:**
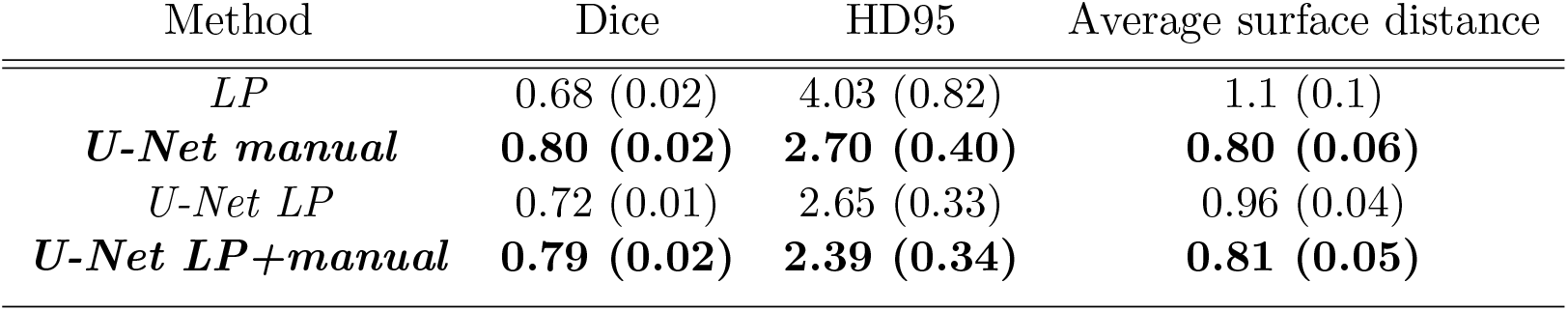
Mean similarity metrics (with standard deviation in brackets) on test set.

These results show that *U-Net LP+manual* achieves the same statistical performance as *U-Net manual* (p-value> 0.05 for all similarity metrics), while outperforming both *U-Net LP* (dice p-value = 6.09 × 10^*-*5^), and *LP* (dice p-value = 1.93 × 10^*-*5^); with the added value that using the propagated labels generates multi-class predictions.

### 4.2. Visual inspection

Fig. 4 depicts an example prediction of each model, showcasing the added value of using a multi-label atlas for multi-class predictions. An important advantage of this multi-label approach over *U-Net manual* is the ability to target small vessels individually. For instance, LSA (white arrow in Fig. 4) remains unsegmented in the *U-Net manual* prediction. Adding multi-class propagated labels enables to detect this vessel.

**Figure 4:**
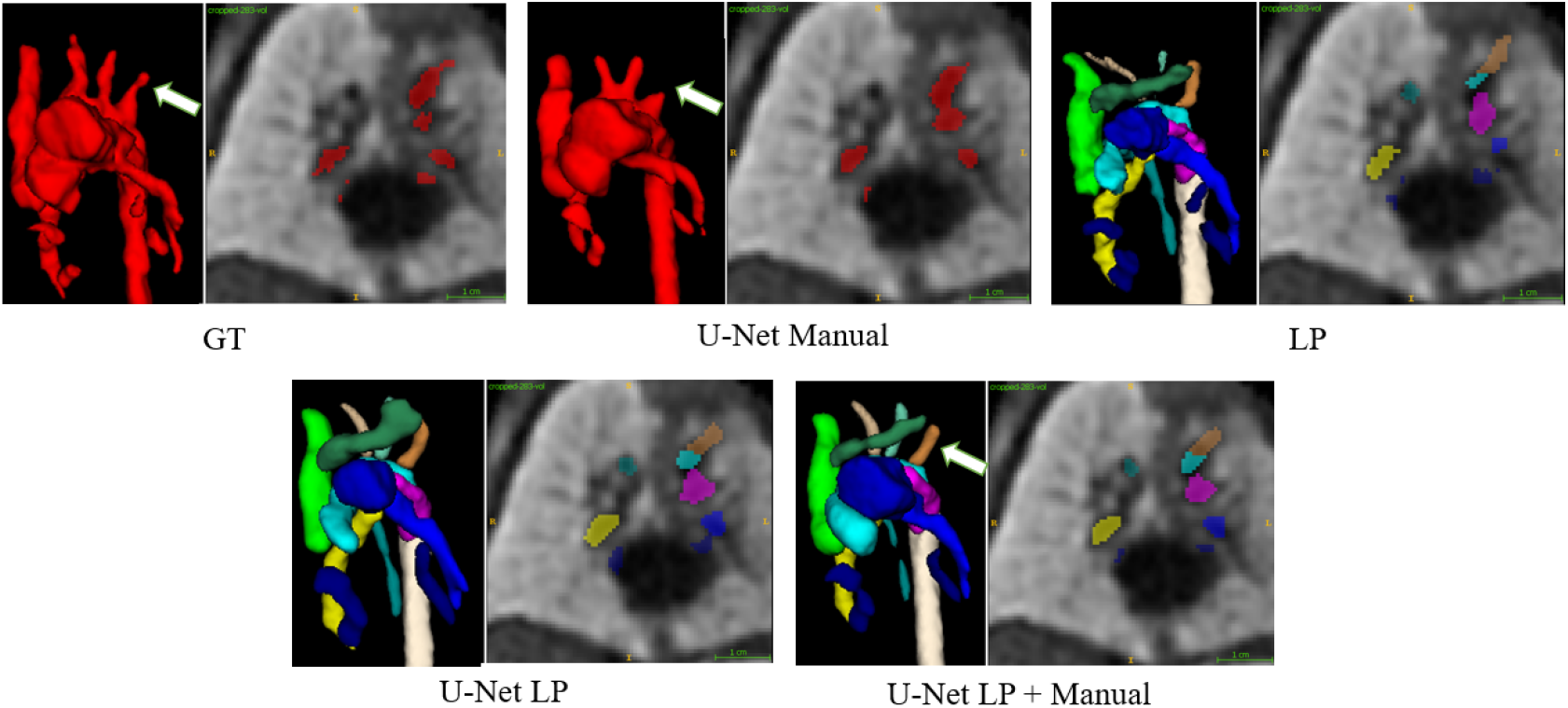
Single and multi-label prediction results for the different experiments. Multi-label frameworks contain two additional vessels: azygos vein and innominate vein (see Fig. 2 for label information).

## 5. Discussion

Our results show that the full framework which combines propagated labels with single-labels leads to synergistic training. Targeting labels individually leads to better segmentation of smaller vessel structures, such as head and neck vessels (see Appendix Section B), while providing multi-class information. Additionally, including a vessels ROI manual label leads to improved overall area segmentation, helping to refine areas of less accurately propagated labels. This can be seen by the improved similarity metric scores (see Table 1) U-Net predictions display compared to *LP* alone.

We achieve a detection rate of 100% for all vessels with *U-Net LP + manual*. Predictions were inspected by a trained clinician, reporting overall optimal results. General comments include partial overestimation of pulmonary arteries and upper pulmonary veins, due to scan quality and the close proximity of distal pulmonary arteries with other respiratory structures. Future work will address this limitation by exploring alternative reconstruction techniques to improve image quality (Uus et al., 2020).

## 6. Conclusion

This study demonstrates the feasiblity of deep learning for multi-label vessel segmentation in black blood T2w reconstructed images, combining label propagation using VoxelMorph with Residual 3D U-Net segmentation. Training experiments explored the combination of single labels of the whole vessels ROI (manual labels) with propagated multi-labels from the atlas, with the inclusion of both labelling information yielding improved predictions. With promising results for subjects with coarctation of the aorta, future work will extend this framework to target other cardiac anomalies, including right aortic arch and double aortic arch.

## Acknowledgments

We would like to acknowledge funding from the EPSRC Centre for Doctoral Training in Smart Medical Imaging (EP/S022104/1).

We thank everyone who was involved in the acquisition and examination of the datasets and all participating mothers. This work was supported by the Rosetrees Trust [A2725], the Wellcome/EPSRC Centre for Medical Engineering at King’s College London [WT 203148/Z/16/Z], the Wellcome Trust and EPSRC IEH award [102431] for the iFIND project, the NIHR Clinical Research Facility (CRF) at Guy’s and St Thomas’ and by the National Institute for Health Research Biomedical Research Centre based at Guy’s and St Thomas’ NHS Foundation Trust and King’s College London. The views expressed are those of the authors and not necessarily those of the NHS, the NIHR or the Department of Health.

## Appendix A. Registration CNN Architecture

Fig. 5 illustrates the CNN architecture employed for label propagation. The input images (atlas and subject) are concatenated along the channel dimension and inputted into the network, with Gaussian smoothing being applied to the resultant velocity field, to ensure diffeomorphism. This is followed by scaling and squaring layers, which yield the final deformation field, used to warp the moving image via a Spatial Transformer Network (STN) (Jaderberg et al., 2015).

**Figure 5:**
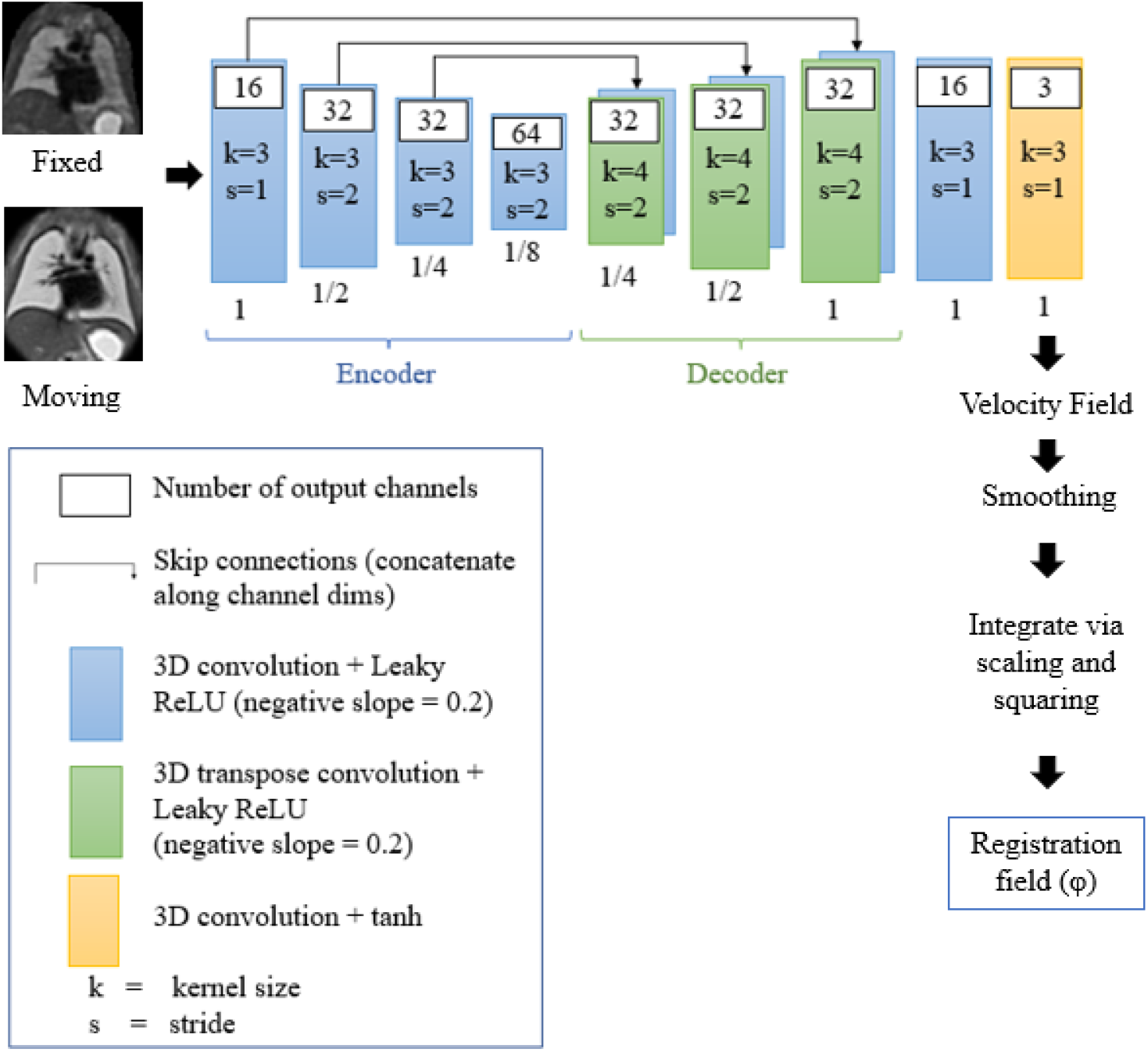
CNN architecture used for registration. The numbers under each convolution representation indicate the volume spatial resolution relative to the input volume. k=kernel size, s=stride.

## Appendix B. Extended Qualitative Evaluation

Here we provide further examples to showcase the advantages of our full multi-label framework (*U-Net LP+manual*) over the alternative models investigated. Fig. 6 showcases an example where *U-Net LP+manual* provides a much more realistic prediction of the shape of the circled vessel (BCA) over *U-Net manual*. The fact that each vessel is targeted individually in our multi-class approach (*U-Net LP+manual*) therefore leads to favourable predictions regarding the accuracy of the shape and structure of small vessels.

**Figure 6:**
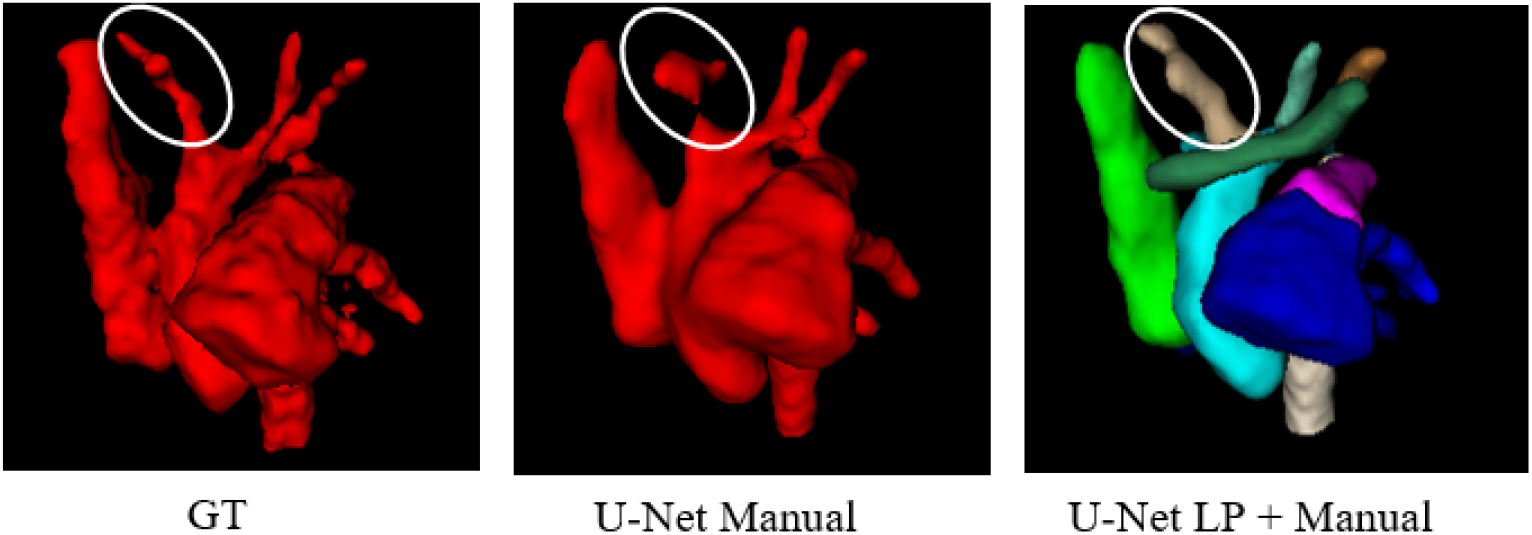
3D Model predictions with BCA circled, showcasing shape prediction improvements with our full framework.

This is further validated by Fig. 7, where *U-Net manual* provides a biologically unrealistic prediction: BCA is joined to SVC (see arrow showcasing joined vessels), in contrast with the prediction from *U-Net LP+manual*. Thus our full framework also proves advantageous for ensuring biologically realistic predictions, again due to vessels being targeted separately.

**Figure 7:**
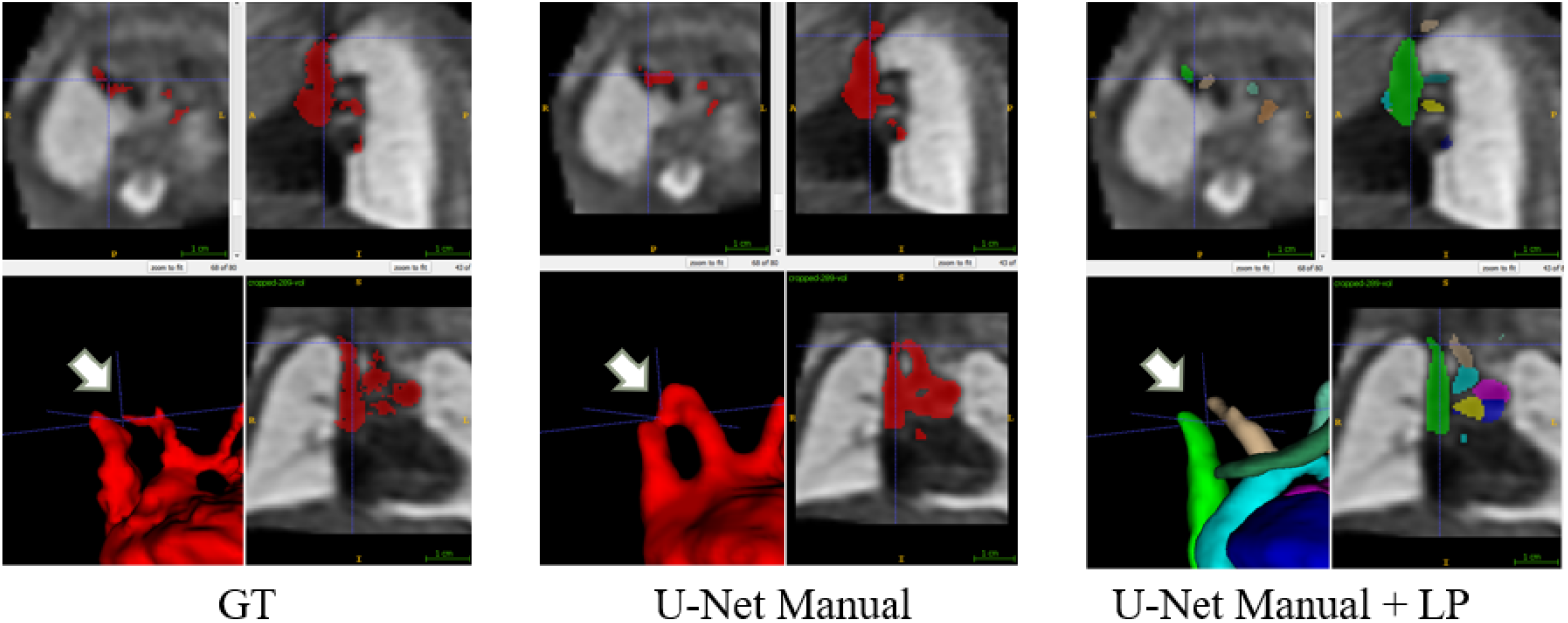
Predictions showcasing inaccurately joined SVC and BCA in the *U-Net manual* prediction (white arrow).

## Footnotes

1. https://gin.g-node.org/SVRTK/

2. https://github.com/Project-MONAI/MONAI/.

